# Neural Evidence of the Strategic Choice Between Working Memory and Episodic Memory in Prospective Remembering

**DOI:** 10.1101/055004

**Authors:** Jarrod A. Lewis-Peacock, Jonathan D. Cohen, Kenneth A. Norman

**Affiliations:** Department of Psychology and Imaging Research Center University of Texas at Austin, Austin, TX 78712, USA; Department of Psychology and Princeton Neuroscience Institute Princeton University, Princeton, NJ 08540, USA

**Keywords:** prospective memory, working memory, episodic memory, fMRI, MVPA

## Abstract

Theories of prospective memory (PM) posit that it can be subserved either by working memory (WM) or episodic memory (EM). Testing and refining these multiprocess theories of PM requires a way of tracking participants’ reliance on WM versus EM. Here we use multi-voxel pattern analysis (MVPA) to derive a trial-by-trial measure of WM use in prospective memory. We manipulated strategy demands by varying the degree of proactive interference (which impairs EM) and the memory load required to perform the secondary task (which impairs WM). For the condition in which participants were pushed to rely more on WM, our MVPA measures showed 1) greater WM use and 2) a trial-by-trial correlation between WM use and PM behavior. Finally, we also showed that MVPA measures of WM use are not redundant with other behavioral measures: in the condition in which participants were pushed more to rely on WM, using neural and behavioral measures together led to better prediction of PM accuracy than either measure on its own.

## Highlights

- We measured participants’ use of working memory for prospective remembering on a trial-by-trial basis using functional MRI.
- Our neural measure of working memory varied according to task conditions that were designed to manipulate strategy use, and it led to better prediction of trial-by-trial prospective memory accuracy than could be achieved based purely on behavioral measures.
- These data provide the strongest connection to date between neural data and prospective memory behavior.

## INTRODUCTION

Prospective memory (PM) refers to our ability to remember to do things in the future. Theories of PM (Cohen and O’Reilly, 1996; Gollwitzer and Brandstätter, 1997; McDaniel and Einstein, 2000) posit that two strategies can be used: Participants can use working memory (WM) to actively monitor the environment for an appropriate time or event (Koechlin and Hyafil, 2007; Gilbert, 2011) or they can store the intention in episodic memory (EM) and hope that it is automatically retrieved when the time comes to act on that intention (McDaniel and Einstein, 2007b; Beck et al., 2014; for related ideas about dual systems involved in PM and control see Cohen and O’Reilly, 1996 and Braver, 2012). PM is typically studied using a dual-task paradigm in which a PM task is embedded in another cognitive task that requires vigilance and frequent behavioral decisions (the “ongoing task”). The PM task requires a response after a particular event (the PM “target”) or after a certain amount of time has elapsed (McDaniel and Einstein, 2007a).

This *multiprocess* view of PM (Cohen and O’Reilly, 1996; McDaniel and Einstein, 2000) raises important questions about when people will rely on one memory strategy vs. the other, and how this strategy choice will affect performance. The current framing of the theory posits an adaptive view of the memory system in which there is a bias to minimize the cognitive demands of the PM task, thereby reducing interference costs from strategic monitoring (Smith, 2003; Einstein et al., 2005; Hicks et al., 2005). Thus an automatic retrieval strategy (relying on EM) is favored whenever possible so as not to overly burden ongoing processing. However, the theory also specifies that some circumstances, when sustained, should favor strategic monitoring (relying on WM); for example, “non-focal” tasks in which identification of a PM target requires attention to features that are not relevant to ongoing processing demands (Einstein et al., 2005; Scullin et al., 2010) and thus might be missed if not actively monitored.

To date, the primary approach to tracking use of strategic monitoring has been indirect: measure RT costs on the *ongoing* task, with the logic being that greater monitoring for the PM target will lead to slower RTs on the ongoing task (Smith, 2003; Einstein et al., 2005; Smith, 2010; Einstein and McDaniel, 2010; Scullin et al., 2010). Neural data has also been used to assist in identifying the strategy in use. fMRI studies of PM have linked strategic monitoring in PM tasks to sustained activity in frontoparietal control networks including anterior regions of the prefrontal cortex (e.g., Reynolds, 2009; McDaniel et al., 2013). In another study, Gilbert (2011) used multi-voxel pattern analysis (MVPA; Lewis-Peacock and Norman, 2014b) of fMRI to successfully decode the contents of WM. However, these measures were unrelated to PM performance. In subsequent analyses, Gilbert et al. (2011) demonstrated that PM accuracy could be predicted by regional increases in fMRI activity and by multivariate measures of similarity between encoding and retrieval. However, most of the above studies used neural measures of WM engagement that were not sensitive enough to predict PM accuracy on a trial-by-trial basis.

One goal of our study was to use a more sensitive, time-varying measure of WM engagement (MVPA decoding of PM target processing) in an effort to improve trial-by-trial predictions of PM behavior beyond what is possible by observing behavior alone. The other goal was to gain a richer understanding of the factors that shape PM strategy use. We designed a PM experiment that manipulated proactive interference and WM load, and that used a non-focal task design – that is, stimuli for the ongoing task (letter strings) that were completely non-overlapping with stimuli for the PM task (faces and scenes). One condition was designed to bias participants to use strategic monitoring (WM*_bias_*; high proactive interference + low memory load), and another was designed to bias participants to rely on automatic retrieval (EM*_bias_*; low proactive interference + high memory load). Using this paradigm, we found that strategic monitoring (measured using MVPA) was both higher overall and more tightly linked to behavior in the WM*_bias_* condition than the EM*_bias_* condition; we also found that our MVPA measure of strategic monitoring improved the ability to predict PM performance from trial to trial, beyond what is possible based on behavior alone.

## MATERIALS AND METHODS

### Participants

Twenty-five participants (14 female; ages 18 to 34, mean = 23.2; all right-handed) were recruited for this study using online scheduling software provided by the Department of Psychology at Princeton University. Participants were compensated with $20 per hour for their participation in the two-hour experiment. Written informed consent was obtained in a manner approved by the Princeton Institutional Review Board.

### Behavioral Paradigm

We developed a task to examine how participants strategically use episodic memory (EM) versus working memory (WM) to remember targets in a dual-task prospective memory (PM) experiment. Participants were shown a series of words while pictures of faces and scenes were presented in the background (Fig. 1a). Participants performed an ongoing task (OG; making lexical decisions about strings of letters) while monitoring for a picture target (a particular face or a particular scene) to reappear. Whereas many studies (see McDaniel and Einstein, 2007a) have used letter stimuli for both the OG task and the PM task, we used pictures (faces and scenes) in the PM task and letters in the OG task (making this a “non-focal” PM task; Einstein et al., 2005; McDaniel et al., 2013). We did this because thoughts about faces and scenes can be tracked effectively using fMRI (Lewis-Peacock & Norman, 2014b); as such, using faces and scenes maximized our ability to use fMRI to track the maintenance of PM targets in WM. Each “PM+OG” trial (in which participants performed both the PM task and the OG task) began with the introduction of a picture target for 2 sec, followed by a 2-sec blank screen, followed by a variable-length sequence of 2-sec memory probes, each containing two pictures and a string of letters. In one-third of the trials, randomly selected, the target introduction screen at the beginning of the trial was blank, indicating to participants that they could ignore all subsequent pictures for the remainder of that trial and focus solely on the OG task (we call these “OG-only” trials). Participants were required to make repeated lexical judgments about the letter strings until the picture target reappeared (between 2 sec and 42 sec after its introduction). In the OG task, a lexical judgment for a given probe required an n-back comparison (n = 1 or 2) of lexical status: i.e., does the current probe have the same lexical status (word or non-word) as the 1-back or 2-back probe? For example, in the 1-back condition, the letter string “apple” (a word) on one probe followed by the letter string “boat” (also a word) on the next probe required a *same* response for the OG task. If, instead of “boat” appearing on the second probe, the letter string “glorb” (a nonword) appeared, the appropriate response on the OG task was *different*. The proportion of same/different responses required was balanced across the experiment. Participants made lexical judgments by pushing a button with the index finger (*same* response) or middle finger (*different* response) of their right hands on a four-button response box. Participants had a 1.9 sec deadline within which to register their responses. For the PM task, participants could identify the picture target when it reappeared by pushing a third button with their pinky finger. Participants were instructed to ignore the OG task on such probes, but they were not prevented from responding to both tasks on any probe (e.g., they could make an OG task response first and then make a PM response, or vice versa, before the response deadline). The PM target reappeared only once per trial, and its reappearance always marked the end of the trial. The probe in which the PM target appeared was varied randomly, from the 1st to the 21st, thus trials varied randomly in their length.

**Figure 1.**
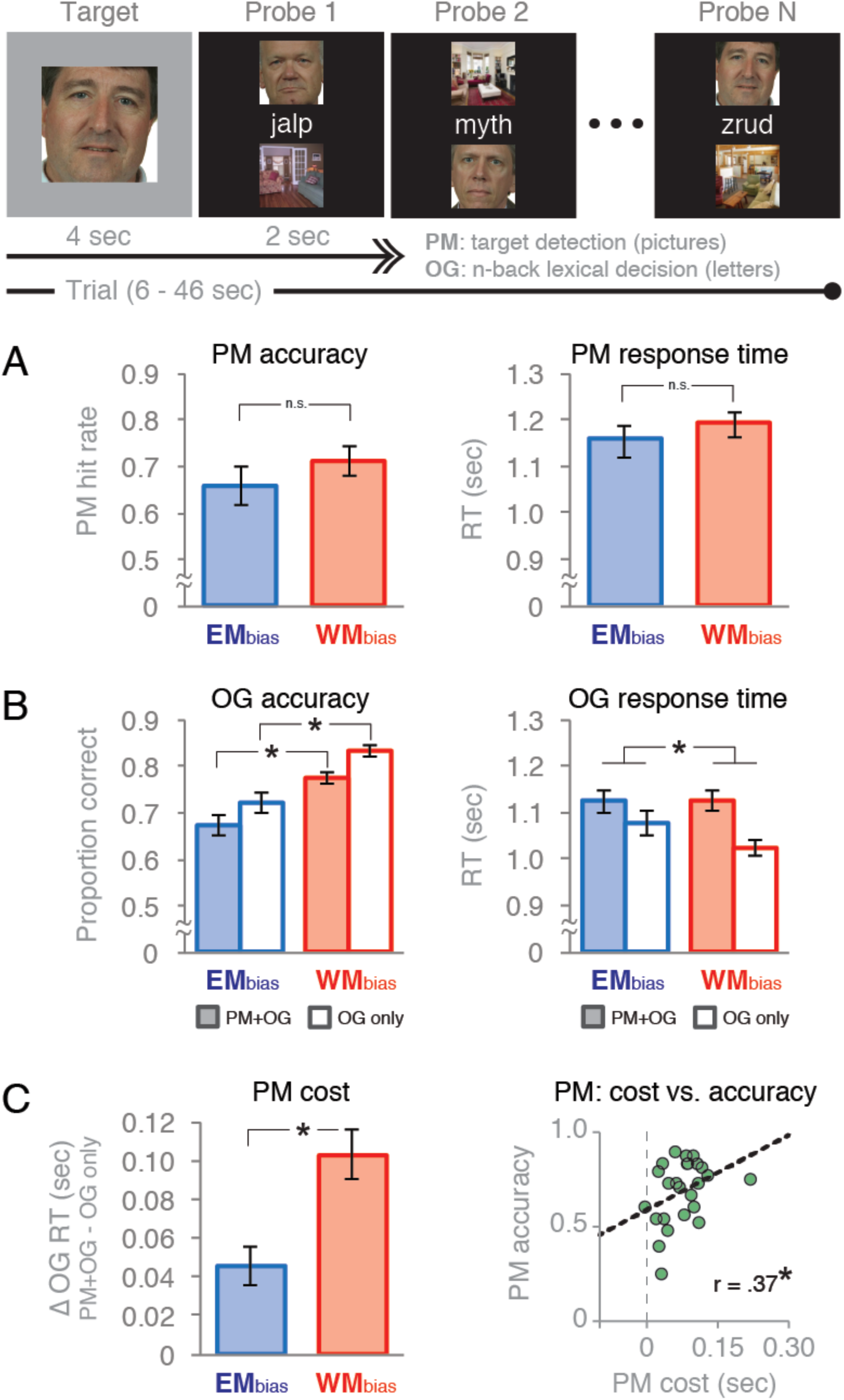
Task diagram and behavioral performance. The dual-task experiment consisted a picture-target detection prospective memory task (“PM”) embedded in an ongoing lexical-decision task (“OG”). Half of the trials were WM*_bias_* trials (1-back lexical decisions and a small set of repeating homogeneous pictures) and half were EM*_bias_* trials (2-back lexical decisions and a large set of trial-unique heterogeneous pictures). Two-thirds of all trials included both tasks (“PM+OG”), and one-third did not require PM responses (“OG only”) Behavioral performance on (A) the PM task in PM+OG trials and (B) the OG task in all trial conditions. (C) Dual-task costs on reaction time in the OG task due to the addition of the PM task (“PM cost”), and its relationship to PM accuracy across participants. Error bars indicate s.e.m.,*p < .05.

Visual feedback was provided after every response. In the OG task, white letter strings immediately turned green if the participant responded correctly, and they turned red if the participant responded incorrectly. In the PM task, if a participant false-alarmed to a distractor picture during a probe (i.e., they incorrectly endorsed a distractor picture as the target picture) the border of the screen turned red for the duration of that probe, but then the trial continued without disruption. When a participant correctly identified a picture target, the border of the screen turned green for the duration of that probe (which was the final one for the trial, as explained above). After this, a screen appeared that indicated whether the participant correctly identified the PM target (black screen with green text stating *“You got it!”*), or failed to identify the target (yellow screen with red text stating *“Oops, you missed it…”*). This feedback was omitted and the screen remained black on OG-only trials. There was a brief 6-sec rest period between each trial to allow for the hemodynamic signal to return to baseline. At the end of each block of trials, participants were shown their average response accuracy for both the OG task and the PM task on that block.

The logic of our experiment was motivated by the multiprocess framework of PM (Cohen and O’Reilly 1996; McDaniel and Einstein, 2000) and the dual mechanisms of control framework (Braver, 2012), which suggest that there are multiple processes than can support prospective remembering: strategic/attention-demanding processes, and also relatively automatic processes. In our PM task, we reasoned that a participant would be able to identify the picture target by either maintaining an active representation of the target (in WM) and strategically monitoring for its reappearance throughout the trial, or spontaneously retrieving the identity of the target (from EM) at the moment that it reappeared. To manipulate participants’ strategy use, we varied the WM load associated with the OG task and the degree of proactive interference associated with the PM targets across trials. Specifically, there were two trial conditions that we refer to as “EM*_bias_*” (high working memory load + low proactive interference) and “WM*_bias_*” (low working memory load + high proactive interference). EM*_bias_* trials were designed to bias participants to use retrieval from EM for prospective remembering. We reasoned that, when a trial involved a higher WM load for the OG task (2-back lexical judgments), participants would be less likely to maintain the picture target in WM, relying instead on retrieval from EM. On these trials, we also used a large set of trial-unique, heterogeneous pictures to reduce the amount of proactive interference amongst the target and distractor pictures and thus further favor use of EM as an effective strategy. Note that, because participants were shown a target only once, they did not have the opportunity to establish a stimulus-response association for that item; therefore, if they were not actively monitoring for the target, we argue that they must have relied on EM to identify it. In contrast, WM*_bias_* trials were designed to bias participants to use WM to maintain the picture target and to actively monitor for its reappearance. We reasoned that high proactive interference (resulting from the repetition of a small, homogeneous set of pictures that repeated within and across trials; Wickens et al., 1963) would interfere with EM retrieval, and that a lower WM load on the ongoing task (Meier and Zimmermann, 2015; 1-back lexical judgments) would encourage a WM strategy for prospective remembering in these trials.

The experiment was divided into six 15-trial blocks of trials, alternating between blocks of WM*_bias_* and EM*_bias_* trials. The trial condition used for the first block was randomly assigned to each participant and counter-balanced across participants. Each block consisted of 12 “real trials” (with exactly one trial of each length, ranging from 10 to 21 probes per trial, inclusive) and three “catch trials” (ranging in length between one and nine probes per trial, with the selection of trial lengths balanced across blocks). Only data from real trials were used for analysis; the catch trials were used to balance cognitive demands throughout the entire trial, i.e., to prevent participants from ignoring the pictures in the first nine probes before engaging in the PM task. There were five trials in each block for each target category (face-target, scene-target, and no-target), consisting of four real trials and one catch trial per category per block. Ignoring catch trials, there were eight PM+OG trials (face or scene target) and four OG-only trials (no target) in each block. The trials were configured such that there were an equal number of probes in each block (62 probes from real trials, and 15 probes from catch trials). Each target category was presented in all of the 12 real-trial lengths and in three of the possible catch-trial lengths in both WM*_bias_* and EM*_bias_* conditions, resulting in a total of 90 trials (72 real trials, 18 catch trials) across the entire experiment. The trials were arranged in this way to reduce participants’ ability to predict the length of any given trial; no participant reported an ability to predict trial length, or knowledge of any structure or pattern of trial lengths across the experiment.

### Stimulus Details

A large collection of face and scene images was gathered through various online and in-house sources. A subset of these stimuli were chosen for this experiment. Words for the lexical comparison task consisted of nouns, verbs, and adjectives selected from an online psycholinguistic database (http://websites.psychology.uwa.edu.au/school/MRCDatabase/uwa_mrc.htm) with concreteness, imageability, and verbal frequency within one standard deviation of the mean of the entire database. Pseudo-words consisted of single-syllable, pronounceable letter strings.

To manipulate proactive interference amongst picture targets, we varied the type and quantity of pictures used in each trial condition. In the WM*_bias_* condition, we used a small set of eight homogeneous face images (adult white males) and eight homogeneous scene images (indoor living rooms) that repeated within and across trials. In the EM*_bias_* condition, we used a large set of heterogeneous faces (789 total; 321 female) and scenes (223 total; 82 indoor) that were trial-unique. The assignment of stimuli to the targets and distractors in each trial was done randomly for each participant.

### fMRI Data Collection

The experiment was presented using Psychophysics Toolbox Version 3 in Matlab running on a Mac Pro. First, we ran a brief scout localizer scan (15 s) to verify that head position was within the designated field of view and to derive automatic AC-PC alignment parameters for subsequent scans. Next, we used a MPRAGE sequence to acquire high-resolution T1-weighted images (TR = 2300 ms, TE = 3.08 ms, 0.9 mm^3^ isotropic voxels, 9 m 0 s acquisition time) while the participants practiced one block of trials in both WM*_bias_* and EM*_bias_* conditions prior to functional scanning. The experiment was divided into six 15-trial blocks of trials (with each block lasting 10 min 3 sec). Total functional scanning time for the experiment was 60 m 18 s. All blocks were preceded by 20 s of dummy pulses to achieve a steady state of tissue magnetization. Between blocks, participants were given a break during which the experimenter checked that the participant was comfortable and alert. Whole-brain images were acquired with a 3T Siemens Skyra MRI scanner. For functional scans, we used a gradient-echo, echo-planar sequence (TR = 2000 ms, TE = 34 ms), with automatic shimming enabled, to acquire T2*-weighted data sensitive to the BOLD signal within a 64 × 64 matrix (196mm FoV, 34 axial slices, 3 mm^3^ isotropic voxels, AC-PC aligned) using integrated parallel acquisition techniques (iPAT) with both retrospective and prospective acquisition motion correction (PACE) enabled.

### fMRI Preprocessing

Preprocessing of the functional data was done with the AFNI (Cox, 1996) software package using the following preprocessing steps (in order): (1) correction for slice time acquisition with 3dTshift, (2) rotation of oblique data to cardinal direction with 3dWarp, (3) resample to a 3 mm^3^ gridset with 3dresample, and (4) realign to the first volume of the Phase 1 data using rigid body alignment with 3dvolreg. Anatomical data were aligned to the first volume of the functional data with align_epi_anat.py. A whole-brain voxel mask was created for each participant by combining the results of 3dAutomask (dilation = 1) across all six functional runs.

### Multi-Voxel Pattern Analysis: Overview

Our goal in analyzing the fMRI data was to sensitively measure processing associated with the PM task. To accomplish this goal, we used multi-voxel pattern analysis (MVPA; Haynes and Rees, 2006; Norman et al., 2006; Lewis-Peacock and Norman, 2014b) to decode face and scene processing (associated with PM task) and lexical decision processing (associated with the OG task) at every time point throughout the trials. The use of category classifiers to track memory maintenance and retrieval has become a standard approach in the memory literature (see Rissman and Wagner, 2012, for a review). We use the approach here to decode the contents of WM, by identifying the degree to which the category of the PM target (a face or a scene) is actively represented prior to its actual reappearance. Neural evidence of such activity could arise from a combination of maintenance of the target (e.g. a particular face) in WM and the processing of distractor pictures from the target’s category (non-target faces) during the trials. Importantly, either source of target-related neural evidence would indicate the use of a WM-dependent strategic monitoring strategy – reactive control relying on EM retrieval should not produce target-related activity prior to the reappearance of the PM target.

### Multi-Voxel Pattern Analysis: Details

fMRI pattern classifiers were trained, separately for each participant, from a subset of all trials and then used to decode independent data from held-out trials (i.e., using k-fold cross validation: training on k−1 blocks of data and testing on the k^th^ block and then rotating and repeating until all blocks had been tested.) Both EM*_bias_* and WM*_bias_* trails were combined for classifier training. Specifically, classifiers were trained on individual brain scans (acquired at 2-sec intervals) from the probe period of each trial, plus data from the 6-sec rest intervals between trials, in the training set. Training scans were labeled according to the category of the picture target from that trial: either *face, scene*, or *no-target*. Scans from the inter-trial intervals were labeled as *rest.* Note that visual input was identical in all three trial conditions (participants were viewing letter strings in the middle of the screen flanked above/below by faces and scenes). The purpose of including the no-target condition in classifier training was to provide additional “negative examples” for the face and scene target classifiers (i.e., trials where faces and scenes were onscreen but participants were not actively monitoring for face or scene targets). As is standard practice in MVPA (Lewis-Peacock and Norman, 2014b) all trial regressors were shifted forward in time by 6 sec to account for hemodynamic lag of the BOLD signal (typically estimated as 4-8 sec to peak after event onset). In each training block, there were 58 scans each for face, scene, and no-target categories, and 45 scans for the rest category, for a total of 290 scans for task categories and 225 scans for the rest category in each training set. We used the trained classifier in each fold of the cross-validation procedure to decode the moment-to-moment cognitive state throughout the held-out block of test data. For each individual 2-sec scan within a test block, the four classifiers (face, scene, no-target, and rest) each produced an estimate (from 0 to 1) of the degree of neural evidence for the condition they were trained to detect.

All pattern classification analyses were performed using the Princeton MVPA Toolbox in Matlab (downloadable from http://www.pni.princeton.edu/mvpa), using L2-penalized logistic regression. The L2 regularization term biases the algorithm to find a solution that minimizes the sum of the squared feature weights. Logistic regression uses a parameter (λ) that determines the impact of the regularization term. To set the penalty λ, we explored how changing the penalty affected our ability to classify the data (using the cross-validation procedure described above). We found that the function relating λ to cross-validation accuracy was relatively flat across a wide range of λ values (spanning from 0.001 to 1,000). We selected a λ value in the middle of this range (λ = 50) and used it for all of our classifier analyses.

### Voxel Selection

To select brain regions to use for the pattern classifiers, we ran a mass-univariate GLM analysis of all functional data using AFNI’s *3dDeconvolve* to identify brain regions that were more strongly engaged during probes (i.e., stimulus displays after the target introduction but prior to its reappearance) on PM+OG trials vs. OG-only trials. This analysis reveals voxels sensitive to the presence of the PM task on top of the OG task. All trial events were modeled with boxcar regressors of appropriate lengths: target (2 sec), probes (2 sec per probe), PM probes (the final probe of the trial in which the target reappears; 2 sec) and feedback (2 sec). A third-order polynomial was used for the null hypothesis, and all basis functions for trial events were normalized to have an amplitude of one. A contrast of *probes from PM+OG trials* > *probes from OG-only trials* was used to calculate percent-signal-change in BOLD data for a second-level group analysis. Only voxels that showed enhanced signal in PM+OG trials were included. The reverse contrast (OG-only > PM+OG trials) revealed a network of voxels, including in the anterior medial prefrontal cortex, that deactivated with the addition of the PM task (Gilbert, 2011; Momennejad & Haynes, 2012, 2013). However, pattern classification of PM stimulus processing (target and distractor pictures) from these regions was at chance levels and therefore these voxels were excluded from further analysis. Participant results in native space were transformed into atlas space and resampled to 4mm^3^ isotropic voxels using AFNI’s *@auto_tlrc* and then spatially blurred with a 8mm FWHM kernel using *3dmerge*. The normalized group data were analyzed using *3dttest++*, and the results were extracted using a cluster radius of four voxels with a minimum cluster size of 40 voxels, and thresholded at the individual voxel level using AFNI’s false discovery rate (FDR) algorithm with q = .05. Finally, this group-level ROI was backward-transformed into each participant’s native space and intersected with that participant’s whole-brain mask to create subject-specific ROIs. The mean number of voxels retained in this “PM-sensitive” mask was 11,686 (SD = 1,122) (Fig. 2a). Finally, a feature selection ANOVA was applied to the preprocessed fMRI data within the PM-sensitive mask to select those voxels whose activity varied significantly (p < .05) between the four categories over the course of the experiment. Feature selection was performed separately for each iteration of the cross-validation classifier training algorithm to avoid any circularity in the analysis (Kriegeskorte et al., 2009). The pattern of activity across these feature-selected voxels was used as the input to the pattern classifiers and the data were analyzed in each participant’s native space.

**Figure 2.**
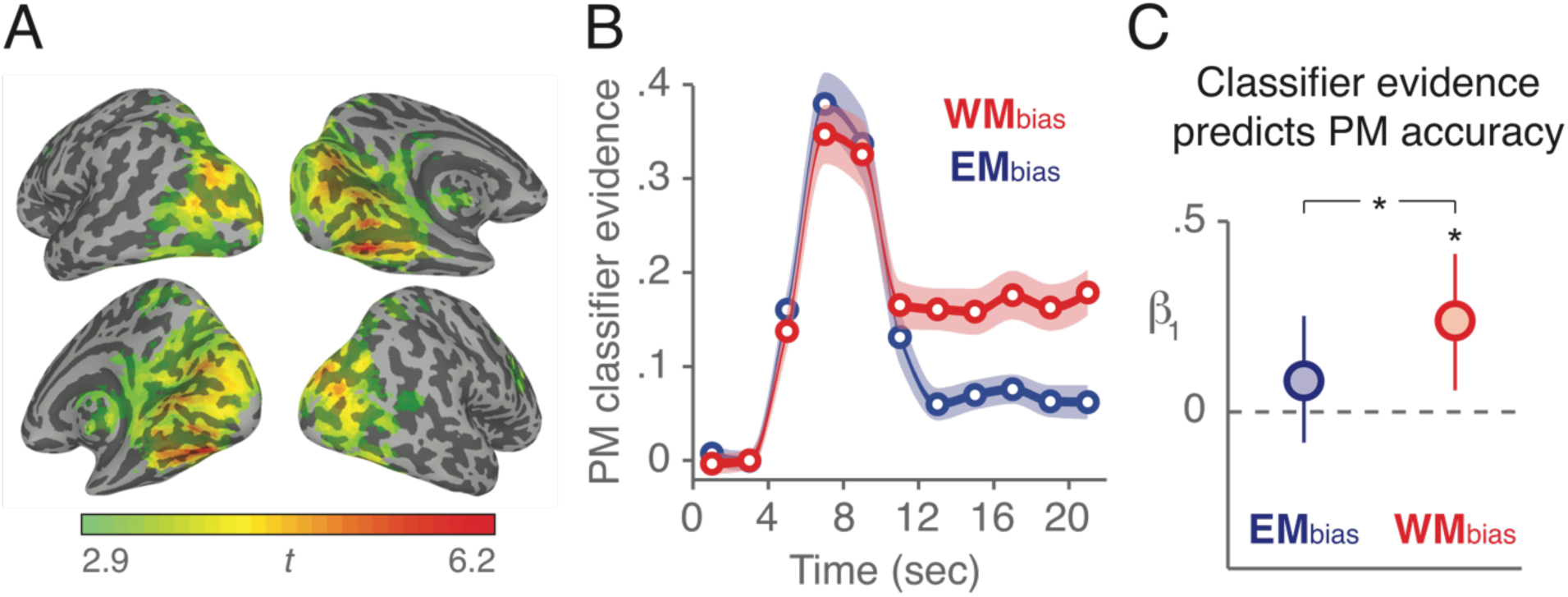
Pattern classification of fMRI data during the delay period predicts PM performance. (A) Voxels that showed significantly greater activity (p < .05, FDR) during probes on PM+OG trials compared to OG-only trials are colored on an inflated atlas brain. This group-level mask was transformed into each participant’s native space and used to mask voxel time series data as input for the pattern classifiers. (B) Trial-averaged classifier evidence for PM trials. PM classifier evidence indicates the difference between target category and distractor category evidence (e.g., “face minus scene” for face-target trials). Error shades indicate +/- 1 s.e.m., interpolated between mean scores from every 2-sec brain scan. Data are not shifted to account for haemodynamic lag. (C) Relating trial-by-trial classifier evidence scores during the delay period (12 to 22 sec) to PM accuracy (hit vs. miss). Data reflect the logistic regression fits (b_1_) between PM classifier evidence and PM accuracy. Error bars indicate 95% bootstrap confidence intervals, *p < .05 for 1,000 bootstrap samples.

### Relating Classifier Evidence to Prospective Remembering

The primary goal of our analysis was to evaluate the relationship between neural classifier evidence for PM monitoring during the trial (prior to target reappearance) to PM accuracy at the end of each trial. We first extracted (separately for each trial in every subject) the levels of face and scene classifier evidence at each time point throughout the trials, and used these data rather than classifier accuracy (whether the correct category had the highest likelihood estimate) for all subsequent analyses (see Lewis-Peacock et al., 2012; Lewis-Peacock and Norman, 2014a). Classifier evidence provides a more sensitive measure of neural processing (and in particular dual-task processing) compared to classifier accuracy because it does not require forced-choice selection of a single “best match” category. To aggregate data across trials that were of varying lengths, we aligned data to the beginning of each trial. Note that the minimum trial length used for analysis contained 10 2-sec probes. Accounting for the target introduction (2 sec) and the brief delay prior to the probes (2 sec), the earliest that the target reappeared in any trial was 2 + 2 + 10*2 = 24 sec. Each trial’s data therefore consisted of 11 brain scans (22-sec, unshifted for hemodynamic lag) aligned to the start of the trial and ending prior to the reappearance of the target.

On a PM+OG trial, target classifier output alone does not show how sensitive the classifier is to the attentional demands of the PM task. High target activation (e.g., “face” on a face-target PM+OG trial) could reflect a highly differentiated attentional state in which target (“face”) is high and distractor (“scene”) is low, or it could reflect a totally undifferentiated state in which both target and distractor are high. Therefore, to track neural processing specifically related to the PM task, we calculated a difference score by subtracting the distractor category evidence from the target category evidence at each time point. Finally, we averaged these difference scores during the PM delay period (t = 12 to 22 secs), which started after the evoked neural response to the target introduction had subsided, and ended before the target reappeared on any of the trials. This method provides a unique neural estimate of PM processing for each trial.

### Statistical Procedures for Assessing Reliability

When analyzing behavioral data (without respect to neural data) and neural data (without respect to behavioral data) we used standard random-effects statistics (paired t-tests, with subjects as a random effect) to assess the reliability of results across participants. For analyses relating neural data to behavior (i.e., PM performance), we combined individual trial data from each participant into a single “supersubject” and subsequently performed all statistical analyses on these amalgamated data (Detre et al., 2013; Kim et al., 2014; Lewis-Peacock and Norman, 2014a), using bootstrap procedures (Efron, 1979) to assess population-level reliability of the results (see details below). We used this approach, chosen *a priori*, instead of the conventional random-effects approach (used elsewhere in the study) in which the *average* results from each subject are used for group-level hypothesis testing. The reason for using the supersubject approach here is that, despite a large amount of imaging data per subject, the total number of behavioral outcomes for each subject was relatively low, making it difficult to reliably estimate the relationship between neural data and behavioral outcomes within individual subjects. Note that each trial lasted between 24 and 46 sec, depending on the number of probes on that trial, but there was only one PM behavioral outcome (hit or miss) on each trial regardless of its length.

In the experiment, each participant (N=25) contributed 12 trials per category/condition combination (e.g., face-target and WM*_bias_* condition) for a total of 300 trials per combination. To assess population-level reliability of the results (i.e., whether or not they driven by a small subset of participants) from each of the analyses, we also ran a bootstrap test in which we resampled data from participants with replacement and re-computed the analyses for this resampled data (Efron, 1979). The population-level reliability of the results was reflected in the proportion of bootstrap samples in which the effect of interest was present.

## RESULTS

### Behavioral Results

#### Prospective Memory Task

We assessed the impact of *trial condition* (WM*_bias_* vs. EM*_bias_*) and *target type* (face vs. scene) on both accuracy and RT in the prospective memory task (PM task; Fig. 1A). With regard to PM accuracy: the hit rate was reliably higher for scene-target trials (p<.01 in both trial conditions), but we nonetheless combined data from both face-target and scene-target trials to increase statistical power for subsequent analyses. PM accuracy was marginally higher for WM*_bias_* trials compared to EM*_bias_* trials (t(24)=1.96, p=.061). There was no interaction between trial condition and target type (F(1,24)=.017, p=.898). The false-alarm rate to non-target items was very close to floor across all trials, although the rate was slightly higher in WM*_bias_* trials than EM*_bias_* trials (0.6% vs. 0.2%; t(24)=3.98, p<.001). Because of this very low false alarm rate, we also calculated PM accuracy using the A’ signal detection metric, which considers both hits and false alarms (Stanislaw and Todorov, 1999). Consistent with the raw hit rates, the A’ signal detection analysis showed a non-significant trend for higher accuracy in WM*_bias_* trials (0.926) compared to EM*_bias_* trials (.914), t(24)=1.71, p=.100. With regard to PM target detection RTs: mean target detection RTs did not differ significantly between EM*_bias_* trials (1.16 sec) and WM*_bias_* trials (1.19 sec), t(24) = 1.33, p=.197. No speed-accuracy tradeoff (i.e., a positive correlation between accuracy and RT) was observed in either condition (WM*_bias_*: r(23) = −.52, p=.008; EM*_bias_*: r(23)= −.08, p=.714).

#### Ongoing Task

We assessed the impact of *trial condition* (WM*_bias_* vs. EM*_bias_*) and *task type* (dual-task: PM+OG vs. single-task: OG only) on both RT and accuracy in the ongoing lexical-judgment task (OG task). There was no main effect of trial condition on OG task RTs (F(1,24)=2.27, p=.15), but participants did respond more slowly on dual-task trials compared to single-task trials (F(1,24)=63.7, p<.001). As noted in the *Introduction*, this slowing of responses in the OG task — a dual-task interference cost that we refer to as “PM cost” — has been interpreted as a behavioral marker for the use of a WM strategy (i.e., that working memory resources were deployed for strategic monitoring of the PM target; McDaniel and Einstein, 2000). We predicted that PM costs would be higher on WM*_bias_* trials, and this prediction was corroborated. There were PM costs in both trial conditions, but PM costs were significantly greater in WM*_bias_* trials (F(1,24)=18, p<.001; Fig. 1C). The same result was obtained when restricting analyses to the PM delay period (t = 12 to 22 secs) that was used to extract neural measurements of PM task processing on each trial (F(1,24) = 16.8, p<.001). With regard to OG task accuracy (Fig. 1B): participants responded more accurately in WM*_bias_* trials compared to EM*_bias_* trials (F(1,24)=56.1, p<.001), and also more accurately on single-task trials compared to dual-task trials (F(1,24)=28.94, p<.001), but there was no significant interaction of trial condition and task type (F(1,24)=0.47, p=.5). These differences in accuracy are consistent with the assumption that the OG task was more demanding in EM*_bias_* trials (2-back) compared to WM*_bias_* trials (1-back); similarly, the greater number of errors in the dual-task condition than the single-task condition is consistent with the greater demands of the former.

#### Individual Differences in PM Performance

PM accuracy and PM cost (i.e., dual-task interference RT costs: OG RT on dual-task trials [PM+OG] minus OG RT on single-task trials [OG only]) both reflect the outcome of strategy choices, and specifically, working memory allocations spread across the dual PM and OG tasks. These two metrics were positively correlated across subjects (r(24) = .37, p = .034; Fig. 1C), indicating that higher PM costs were associated with better PM performance. This relationship was previously reported by Smith (2003; but see McNerney and West, 2007). The correlation was significant for WM*_bias_* trials (r(24) = .45, p = .024), but not for EM*_bias_* trials (r(24) =.199, p = .340). However, the correlation did not significantly differ between the two conditions (z = −.755, p=.45).

### fMRI Results

#### Classifier Cross-Validation

A univariate GLM was used to identify voxels that were more active on PM+OG trials vs. OG-only trials (see Methods). These voxels were located mostly in ventral temporal, occipital, and parietal areas (Fig. 2a), and were used as input for pattern classification. Pattern classifiers, trained and tested separately for each participant, successfully distinguished task-related brain activity on (1) face-target trials, (2) scene-target trials, and (3) no-target trials, and also task-unrelated brain activity during (4) rest periods between trials. Cross-validated classifier accuracy was greater than chance-level performance (0.25) for all four categories (all p’s < .001), and all three task-related categories showed higher accuracy in WM*_bias_* trials vs. EM*_bias_* trials (all p’s < .001), while accuracy for rest-period activity did not differ between conditions (t(24)=0.189, p=0.852). Classification performance did not differ between face-target and scene-target trials (p>.4), therefore classifier estimates from all target trials were relabeled and combined (e.g., on a face-target trial, the “face-target” classifier’s output was relabeled as “target” and the “scene-target” classifier’s output was relabeled as “distractor”).

The PM classifier evidence scores (i.e., “target - distractor”) showed a significant interaction of *trial condition* (WM*_bias_* vs. EM*_bias_*) x *time* (target introduction: 4-12 secs vs. delay period: 12-22 secs; F(1,24) = 32.37, p < .001). PM evidence did not differ between WM*_bias_* and EM*_bias_* trials during the early part of the trials when the target was introduced and encoded into WM (8 secs following target introduction, t = 4 to 12 secs, p = .598), but it did differ during the subsequent delay period (F(1,24) = 31.26, p < .001; Fig. 2b) with higher PM evidence on WM*_bias_* trials during the delay. The fact that classifier performance was matched for WM*_bias_* and EM*_bias_* trials during the early (encoding) phase of the trial suggests that subsequent differences in classification cannot be attributed to generally poorer classification of the large set of heterogeneous face and scene stimuli on EM*_bias_* trials vs. the small set of homogeneous stimuli on WM*_bias_* trials; rather, the difference in classifier performance appears to be specific to the delay period and likely reflects a greater use of WM for the PM task on WM*_bias_* trials relative to EM*_bias_* trials.

It is possible, however, that lower classifier performance in EM*_bias_* trials was due to increased measurement noise in that condition, resulting from the presence of a more demanding OG task (2-back). To address this possibility, we computed the within-trial variability (standard deviation) of classifier evidence scores during the delay period. Variability of target evidence was higher in EM*_bias_* trials relative to WM*_bias_* trials (0.123 vs. 0.100; t(24)=3.5, p<.001), however, distractor evidence was *less variable* in EM*_bias_* trials relative to WM*_bias_* trials (0.141 vs. 0.151; t(24)=2.1, p=.024), which by itself is inconsistent with a “measurement noise” account of these neural data. However, we also assessed the effects of extra “measurement noise” by adding random noise, sampled from a Gaussian distribution (mu=0, sigma=0.100), into both the target and distractor classifier evidence scores from WM*_bias_* trials, thus simulating a “noisy WM*_bias_*” condition. If WM*_bias_* vs. EM*_bias_* differences were merely due to extra measurement noise in the latter, then the qualitative pattern of results in the “noisy WM*_bias_*” condition should match the pattern that was observed in the EM*_bias_* condition; conversely, if the results do not match, this indicates that additional measurement noise alone can not account for WM*_bias_* vs. EM*_bias_* differences. Adding noise in this fashion increased the variability of both target and distractor measurements in WM*_bias_* trials (ps<.001), and also increased their variabilities relative to EM*_bias_* trials (ps < .004). However, the added noise did not change the mean evidence for either target or distractor (both ps > .178). The mean distractor evidence remained higher in EM*_bias_* trials relative to the “noisy WM*_bias_*” trials (0.699 vs. 0.653; t(24)=3.4, p=.001), and most importantly the PM evidence (target – distractor evidence) remained lower in EM*_bias_* trials (0.066 vs. 0.166; t(24)=6.2, p<.001).

### Relating Classifier Evidence to PM Performance

For each trial, we calculated a PM classifier evidence score (as described above) and used this score to predict PM performance (hit or miss) at the end of each trial. Logistic regression was used to relate each continuous classifier evidence score to the binary outcome variable of PM accuracy. To increase statistical power for this regression, individual trial data were combined across subjects into a supersubject analysis (see *Methods*), and reliability of the regression analysis was assessed using a bootstrapping procedure.

Consistent with the prediction that participants would rely more heavily on WM in the WM*_bias_* condition, PM classifier evidence scores were positively correlated with PM accuracy in WM*_bias_* trials (logistic regression β_1_ > 0 in 99.6% of 1,000 bootstraps), but they were not reliably correlated with PM accuracy in EM*_bias_* trials. Regression coefficients were higher for WM*_bias_* trials compared to EM*_bias_* trials in 94.4% of bootstraps (Fig. 2d), indicating that trial-by-trial fluctuations in PM classifier evidence were more predictive of behavior on WM*_bias_* trials compared to EM*_bias_* trials.

### Relating OG Task Behavior to PM Performance

As has previously been observed, OG task behavioral metrics (accuracy and RT) were also predictive of PM accuracy across trials. OG task accuracy was positively correlated with PM accuracy in both trial conditions (β_1_ = .284, which was positive on 99.3% of bootstraps in both trial conditions), with no reliable difference between the coefficients in the two conditions. OG task RT was weakly, but positively correlated with PM accuracy in both trial conditions (β_1_ = .212 for WM*_bias_* and β_1_ =.160 for EM*_bias_*). These coefficients were positive on 93.2% and 89.1% of bootstraps, respectively, with no reliable difference between the coefficients in the two conditions.

### Combining Behavioral Data and Neural Data to Predict PM Performance

The findings above indicate that both behavioral and neural measures were predictive of PM performance from trial to trial. Here we address the question of whether neural evidence provided extra predictive power beyond what what was possible from behavioral observations alone. Using our neural measure (PM evidence) and the two behavioral measures (OG accuracy and RT) together in a three-predictor logistic regression model explained the most variance in PM accuracy scores. To control for differences across models in the number of predictors, we used a leave-one-participant out cross-validation procedure (Hastie et al., 2005): each model was fit using data from N-1 participants and then used to predict data from the held out participant. Average log likelihood values across all iterations for each model were used to calculate Bayes factors (B_10_), which assess the relative likelihood of each model (Kass and Raftery, 1995) taking into account the number of predictors. The three-predictor model outperformed the two-predictor model (OG accuracy and OG RT) in WM*_bias_* trials (log_10_(B_10_) = 3.12; this constitutes “decisive” evidence according to Kass and Raftery, 1995), but not in EM*_bias_* trials (log_10_(B_10_) = 0.38; this is “not worth more than a bare mention” according to Kass and Raftery, 1995). Importantly, this analysis demonstrates that the neural measurements of PM task processing contributed predictive power concerning PM performance on a given trial, above and beyond what could be predicted based on observable OG task behavior alone.

### Individual Differences in Relating Neural Measurements to Performance

The neural findings also revealed individual differences in performance across participants. The amount of PM classifier evidence for a given participant was positively correlated with both overall PM accuracy (r(25)=.73, p<.001) and overall dual-task PM costs (r(25)=.43, p<.05; Fig. 3).

**Figure 3.**
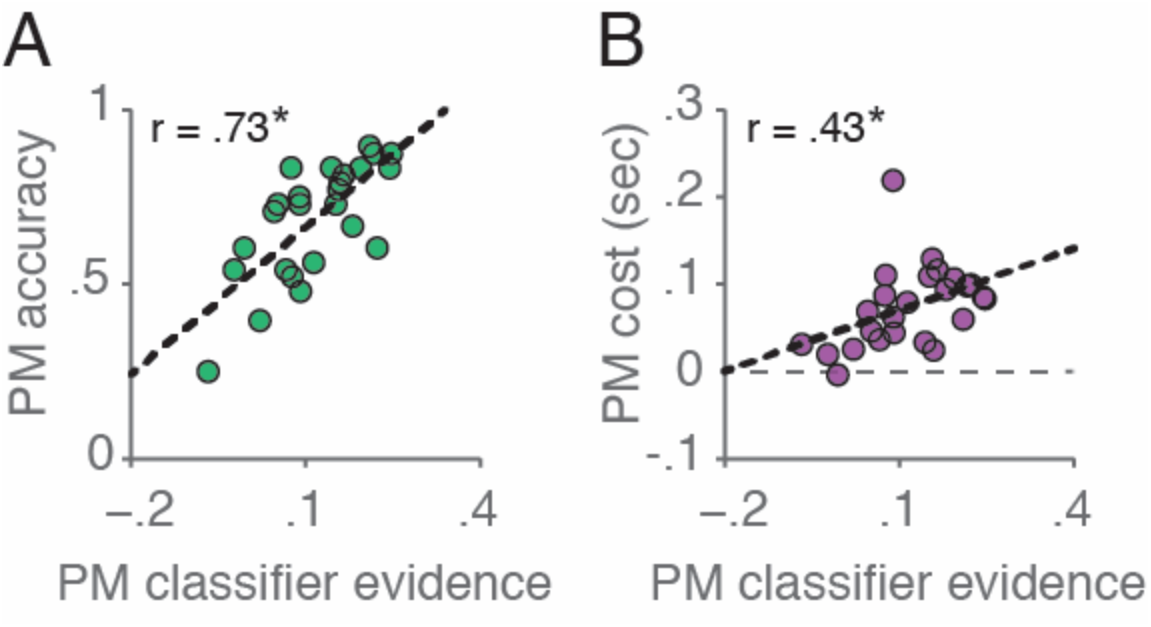
Classifier evidence scores predict PM accuracy and dual-task PM costs across participants. Higher PM classifier evidence was predictive of (A) better PM accuracy and (B) higher dual-task costs. *p < .05.

## DISCUSSION

We developed a novel experimental paradigm, designed to bias strategy choice for prospective memory (PM) on a trial to trial basis, by concurrently manipulating proactive interference and working memory load. When participants were biased to use working memory (WM) over of episodic memory (EM), the PM task exerted a larger cost on the ongoing task (as evidenced by slower RTs) — this dual-task interference cost is considered to be a behavioral hallmark of strategic monitoring (Smith, 2003; Einstein et al., 2005; Scullin et al., 2010; Meier and Zimmermann, 2015). Previously, behavioral interference costs have been used to demonstrate effects of a wide range of factors on PM strategy use, including: the availability of cognitive resources and the sensitivity to interference costs (Marsh et al., 2003; Smith, 2003; Marsh et al., 2006), the instructional emphasis on the PM task and the duration of the ongoing task (Einstein et al., 2005), the degree & type of planning (Mäntylä, 1996; Burgess and Shallice, 1997), and individual differences in cognitive capacities and personality characteristics (McDaniel and Einstein, 2000). In this study we identified another set of task demands that bias participants towards a WM strategy: *high proactive interference* (which makes it harder to use EM) combined with a *low working memory load* (which makes it easier to use WM).

In addition to behavioral evidence, we used pattern classifiers applied to fMRI data from visual processing regions in temporal and occipital cortices during the PM task to acquire second-by-second readouts of neural activity associated with the use of WM to maintain and/or monitor for a PM picture target. Neural readouts of PM processing were higher when participants were biased to use WM (vs. EM). Furthermore, across trials, PM classifier evidence was more predictive of successful PM performance when participants were biased to use WM compared to when participants where biased to use EM, even though PM accuracy was equivalent across the two conditions. These findings complement and extend prior work that has leveraged fMRI data to dissociate PM strategies using activity from a distributed network of brain regions. For example, work by McDaniel et al. (McDaniel et al., 2013) showed that activity in frontoparietal control networks was greater in conditions that require greater levels of strategic monitoring (e.g., non-focal vs. focal PM targets; for other relevant data, see Reynolds et al., 2009; Burgess et al., 2011; McDaniel et al., 2013; Barban et al., 2014; Beck et al., 2014).

Our interpretation of the converging behavioral and neural data of this study is that participants used WM more for the PM task during trials when they were biased to do so. A plausible alternative explanation for the neural findings, however, is that PM evidence (i.e., target evidence – distractor evidence) was lower in EM*_bias_* trials due to increased measurement noise, resulting from the presence of a more demanding OG task (2-back) in that condition. To address this concern, we assessed the variability in classifier readouts in both conditions, and then simulated the effects of adding measurement noise to the classifier in WMbias trials. Together, the observations in EM*_bias_* trials of (a) more stable distractor evidence relative to WM*_bias_* trials, and (b) lower PM evidence compared to simulated “noisy WM*_bias_*” trials are incompatible with the possibility that differences in classifier performance between the two conditions could be due to increased measurement noise in EM*_bias_* trials. While alternative explanations may exist for any individual portion of these data, the idea of greater WM use in the WM*_bias_* vs. EM*_bias_* condition parsimoniously explains, with a single mechanism, the full set of neural and behavioral findings, including the correlations between these measures.

Crucially, our neural measure of WM use provided additional predictive power concerning PM performance, beyond that provided by behavior alone. This demonstrates how decoding the contents of WM from fMRI data can provide unique evidence concerning the selection and success of cognitive strategies deployed during complex cognitive tasks. Prior work has shown that the content of delayed intentions (e.g., waiting for a word versus a picture to reappear) can be decoded from posterior cortical regions (Gilbert, 2011). However, these neural measurements were unrelated to behavioral metrics of PM performance (but see Gilbert et al., 2011). Here, we found that (particularly when participants were biased to use WM to store their delayed intention) neural readouts of WM use were diagnostic of PM target detection accuracy on a trial-by-trial basis. Across participants, these neural measures were also diagnostic of individual differences both in PM accuracy and dual-task interference costs.

It is important to note that both the EM*_bias_* and WM*_bias_* conditions in our experiment were “non-focal” tests of PM, insofar as the stimuli pertaining to the ongoing task (letter strings) were distinct from the stimuli that pertained to the PM task (faces and scenes). As such, both conditions required some degree of strategic monitoring: Specifically, participants had to allocate some attention to the stream of faces and scenes, in order to be able to detect the face or scene target when it appeared. Our key prediction was that, in the EM*_bias_* condition, participants might favor monitoring for the target *category* without actively maintaining the *specific identity* of the target stimulus. For example, if the participant knew that the target was a face, they might actively monitor the stream of faces, with the expectation that the target face would trigger episodic retrieval of its status as a target. By contrast, in the WM*_bias_* condition, participants might devote extra WM resources to monitoring for the specific target face. The fact that *some* strategic monitoring was required in both conditions fits with the finding that dual-task costs (on OG task reaction times) were obtained in both conditions, although (as predicted) they were larger in the WM*_bias_* condition.

Our data are consistent with the multiprocess view of PM (Cohen and O’Reilly, 1996; McDaniel and Einstein, 2000) and the dual mechanisms view of PM (Braver, 2012), which posit distinct routes for successful PM performance: proactive control via working memory, and reactive control via episodic memory. Here, we found behavioral and neural signatures of the former, but unlike previous work (Braver et al., 2003; Reynolds et al., 2009; McDaniel et al., 2013), we did not find reliable neural signatures for the latter. The reason could be methodological, insofar as the design of our experiment was tailored to identify sustained working memory processing during the PM task (long delay periods, and therefore relatively few PM trials). This may have reduced our ability to detect transient activity spikes associated with episodic memory retrieval in the resulting small set of PM trials. Regardless, we found that PM performance was preserved for trials in which our neural measurement of WM processing was low (e.g., EM*_bias_* trials), and thus PM performance had to have been supported by some process that complemented the diminished engagement of active monitoring via WM. The multiprocess view of PM suggest that it is a reactive control process by which the delayed intention is encoded into episodic memory as a stimulus/response association (e.g., “when this picture appears, hit a special button”) and the retrieval of this intention is automatically triggered by the reappearance of the stimulus.

One potential limitation of the MVPA measure we used to index WM engagement is that it is sensitive to *both* of the types of monitoring described above: checking the “stream” of face stimuli (without holding a specific face in mind), and monitoring for a specific face. There is no way to disentangle the contribution of these two processes to our neural measures. Nevertheless, both sources reflect the engagement of some form of strategic monitoring. The finding that our MVPA measure of WM function was stronger in WM*_bias_* trials is consistent with the engagement of *both* monitoring processes on those trials (checking the target category “stream”, plus monitoring for specific target stimuli), whereas only the checking process may have been engaged in EM*_bias_* trials. This might also explain why our neural measure of WM was more predictive of behavior in WM*_bias_* trials: monitoring for the specific target stimulus should substantially increase the likelihood of responding correctly when the target appears, thus WM use should be correlated with correct responding. By contrast, merely checking the target category stream (without actively holding the correct stimulus in mind) is insufficient to ensure a correct response – even when the target stimulus is seen, it might fail to trigger the corresponding episodic memory, resulting in a PM error; thus, there should be a weaker relationship between WM use and correct responding.

In conclusion, we designed an experiment to bias participants to use either WM or EM to solve a PM task while simultaneously engaged in a demanding ongoing task. Using MVPA to measure strategic monitoring (Lewis-Peacock and Norman, 2014b), we validated that our manipulation was effective in biasing participants’ strategies. More generally, using MVPA improved our sensitivity to detect participants’ strategy use beyond what was possible based on behavior alone, leading to improved trial-by-trial predictions of PM accuracy. Future work can leverage these improvements to further characterize the factors that shape PM performance both within and across individuals.

## Conflict of Interest

none

## Acknowledgements

We thank Matthew Salesi and Vivian DeWoskin for assistance with behavioral piloting of this experiment. We also thank Jeremy Manning, Jordan Poppenk, Nicholas Turk-Browne, Ida Momennejad, Richard Lewis, and Satinder Singh for many helpful discussions during the development and analysis of this experiment. This research was funded by NIMH grant R01 MH069456 awarded to K.A.N. and by through the support of a grant from the John Templeton Foundation. The opinions expressed in this paper are those of the authors and do not necessarily reflect the views of the John Templeton Foundation.

